## Supplementary information and figures for "Weak parent-of-origin expression bias: Is this imprinting?"

#### Comparison of four studies chosen for analysis

For best comparison, we ignored all non-mouse data, as well as imprinting data from the mouse X chromosome, which was only analysed in one of the studies[17] as well as any non-imprinting work performed. Crowley et al. used CastEiJ, PWK/PhJ, and WSB/EiJ mice in reciprocal crosses of all combinations whereas the other three studies all used C57BL/6 x CastEiJ reciprocal mouse crosses (Supplementary Table S1)[16–19]. As RNA-seq starting material, Perez and colleagues used mouse cerebellum at postnatal day (P)8 and P60 (Dataset C) [19]. Bonthuis *et al.* used mouse liver and skeletal muscle, as well as two highly specific brain regions, the arcuate nucleus (ARN) of the hypothalamus and the dorsal raphe nucleus (DRN) of the midbrain, at 8-10 weeks of age (Dataset B) [17]. Crowley and colleagues used whole brain at P23 for their imprinting analysis (Dataset D)[18]. The Babak *et al* study generated an imprinting atlas from 23 tissues from between 35- and 45-days' post-partum (including whole brain and six brain regions) and three fetal tissues, in addition to reanalysing data from five previous studies (Dataset A)[16]. Details of the four different studies can be found in Supplemental Table S1.

Although all studies used the same methods for RNA-seq (Illumina Tru-Seq RNA Kit v2), each group used a different number of replicates. The Crowley and Perez studies used 6 males and 6 females for each reciprocal cross in each tissue. Whereas Bonthuis and colleagues used 8 or 9 replicate females for each cross and tissue. Babak *et al*. pooled tissues dissected from 2 males for some tissues so only one library was generated for each reciprocal cross in most cases. The Bonthuis and Perez studies generated 59 bp single-end data and Babak et al. and Crowley et al. generated 90 bp and 100 bp paired-end data respectively.

All four studies also used different statistical analysis to call allele specific expression. Babak *et al.* estimated ASE from the cumulative binomial distribution with the random expectation set to 50% (equal expression from both alleles). The log10 of the least significant p value between the two reciprocally crossed tissue is taken as the imprinting score with the paternal bias set as negative and maternal expression as positive. The Bonthuis study used generalized linear modelling with false discovery rate (FDR) estimation centred upon a permutation-based approach. For each data set they used a p value that yielded an FDR of 1%. Crowley *et al.* used the method detailed in Zou *et al.* 2014 to jointly model total number of reads (TReC) and number of allele-specific reads (ASE) of each gene, which, they claim, significantly boosts power for detecting strain and particularly parent-of-origin effects. Finally, the Perez study used an in-house pipeline, Bayesian regression allelic imbalance model (BRAIM), which accounts for all sources of variability in the experimental design i.e. cross, sex and age.

A critical analysis of using RNA-seq to infer imprinted expression highlighted the need for validation via an independent method [15]. However, only three of the studies included such validation. One of the studies (Perez et al., 2015) conducted a full quantitative validation by pyrosequencing of all novel genes. They found nine false positives, which are not included in our meta-analysis. Bonthuis and colleagues validated only a subset of 18 novel genes by pyrosequencing, of which 8 failed[17]. Since this approach was not very systematic, testing only 12% of all novel genes, we included the original RNA-seq results including the false positives in our meta-analysis. Babak and colleagues validated most of the novel genes by pyrosequencing but in only one of all respective positive tissues[16]. They only made RNA-seq data available for novel genes that either passed validation (13 genes) or could not be tested (12 genes), so we could not include other results. The final numbers of ASE autosomal genes reported for each study were as follows: Babak *et al.* - 125 (Dataset A), Bonthuis *et al.* - 210 (Dataset B), Perez *et al.* - 115 (Dataset C) and Crowley *et al.* – 95 (Dataset D) [16–19].

### Supplementary Figures

### Supplementary Figure S1

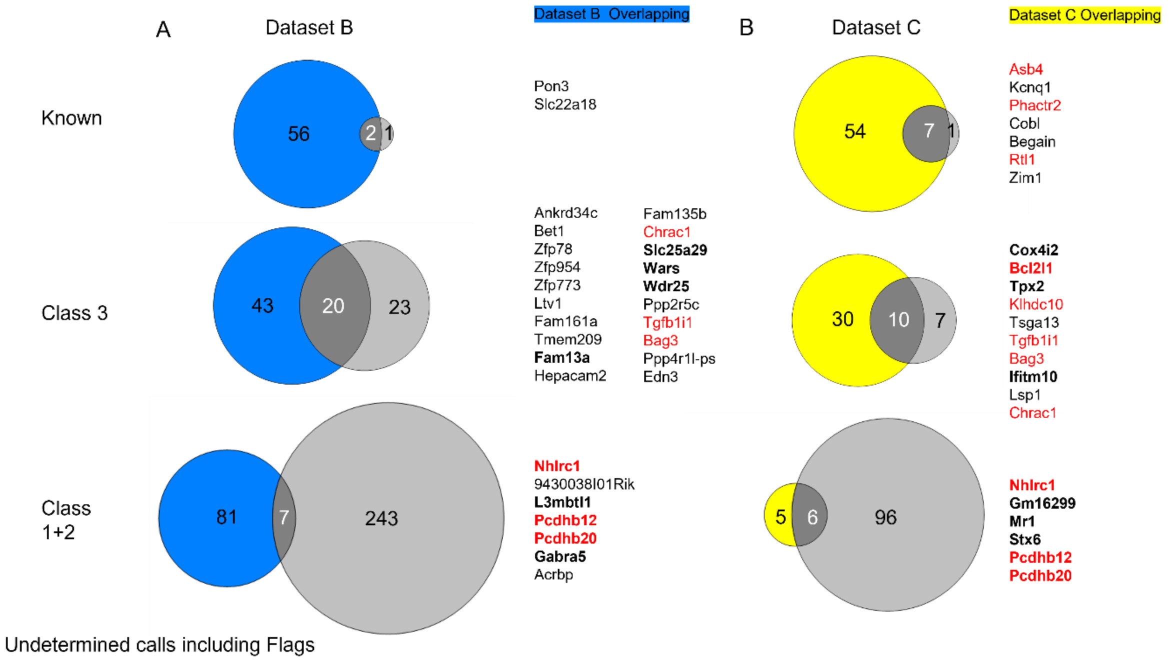


Figure S1 – Overlapping underdetermined genes called between Dataset B (A) and Dataset C (B) and ISoLDE pipeline. Number of ISoLDE called genes are in pale grey, overlaps with original study are in dark grey. Overlapping genes are listed to the right. Gene in red were flagged by ISoLDE and genes in bold were validated by pyrosequencing.

**Supplementary Figure S2**

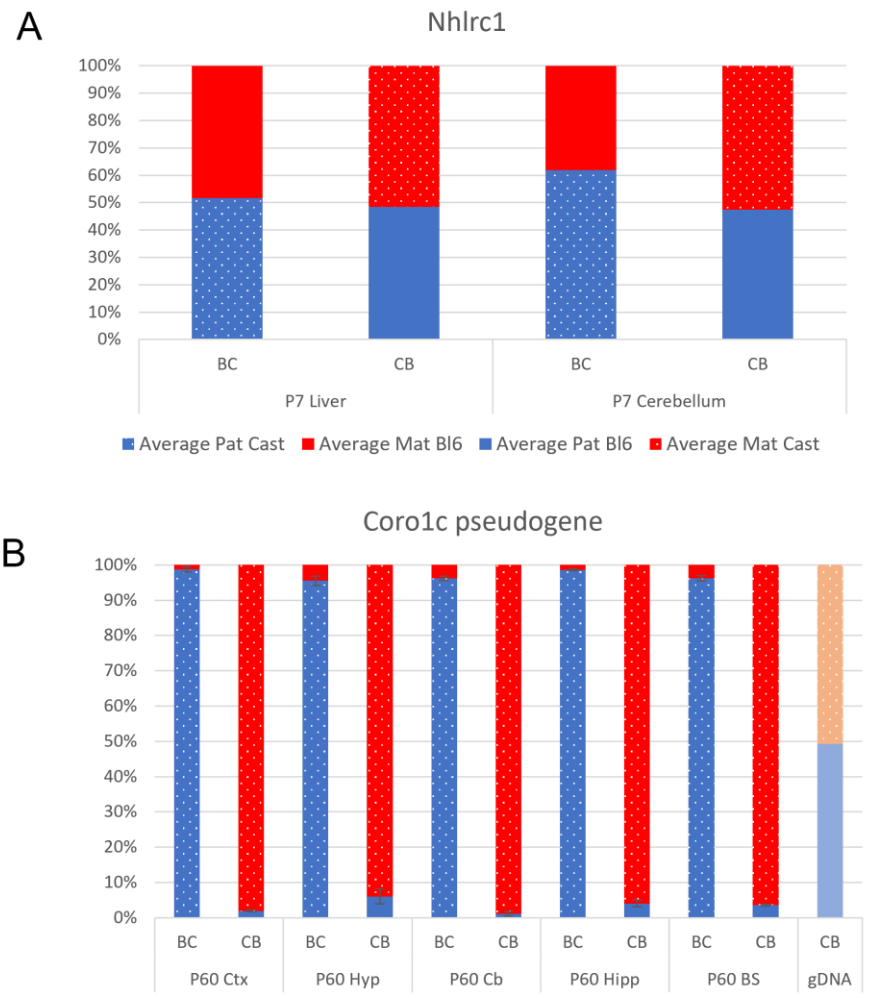


Figure S2 – (A) *Nhlrc1* expression bias in P6 cerebellum and liver which were also used for bisulfite sequencing analysis. (B) Allelic balance of expression of *Coro1c* pseudogene that overlaps the *Nhlrc1* DMR. Graph shows mean expression (%) from the paternal allele (deep blue) and maternal allele (red). C57BL/6 x CastEiJ (BC) and CastEiJ x C57BL/6 (CB) crosses. Castaneus allele is denoted by spotted pattern. Standard error of the mean is shown, N = 2 for *Nhlrc1* and 3 for *Coro1c* pseudogene.

**Supplementary Figure S3**

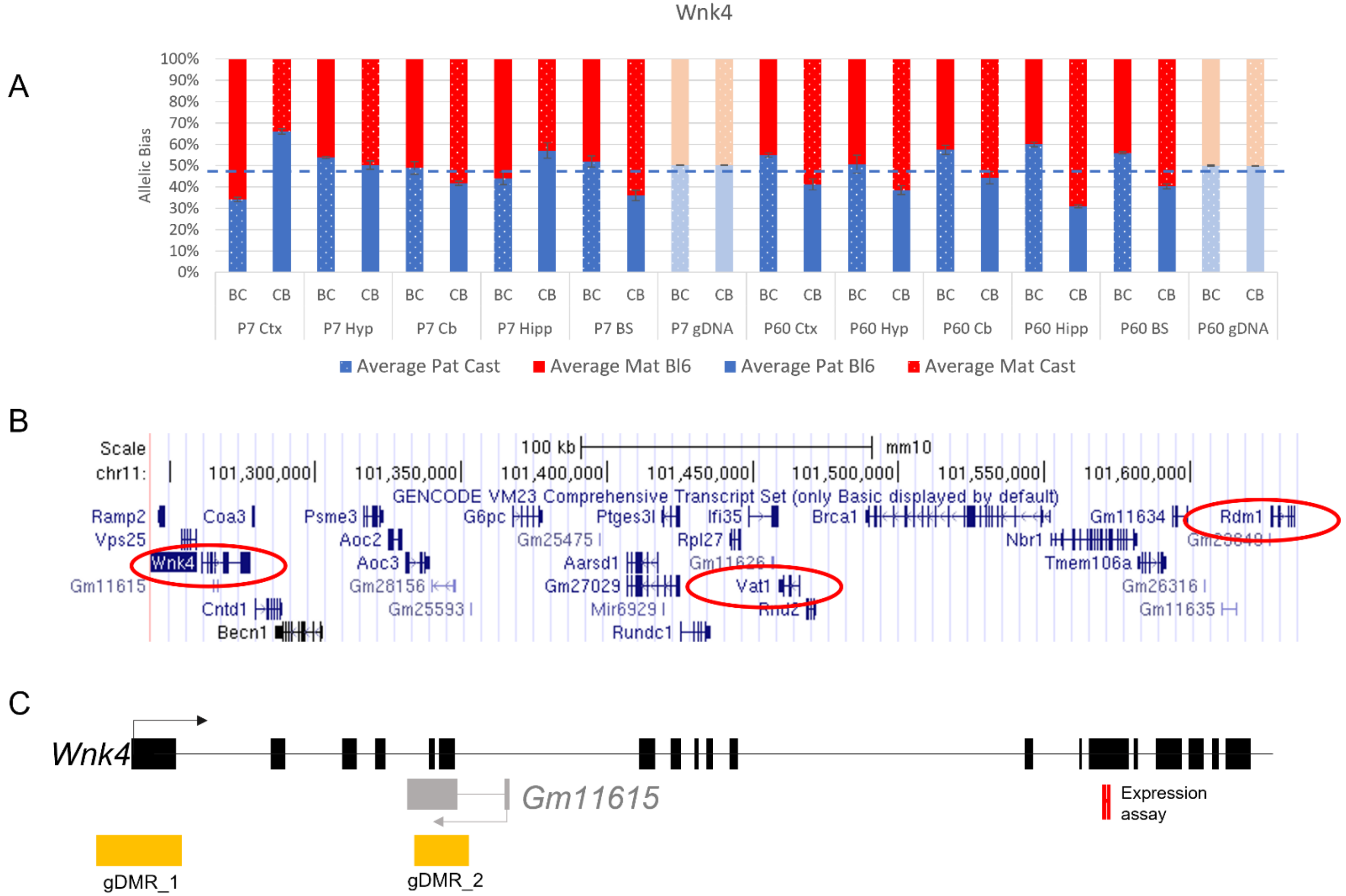


Figure S3 – (A) Weak bias in *Wnk4* gene. (B) location of genes in novel cluster, (C) location of gDMRs and antisense transcript. Graph shows mean expression (%) from the paternal allele (deep blue) and maternal allele (red) C57BL/6 x CastEiJ (BC) and 4 CastEiJ x C57BL/6 (CB) crosses. Castaneus allele is denoted by spotted pattern. Standard error of the mean is shown, N = 3 or 4. Data are normalised for amplification bias in gDNA.

**Supplementary Figure S4**

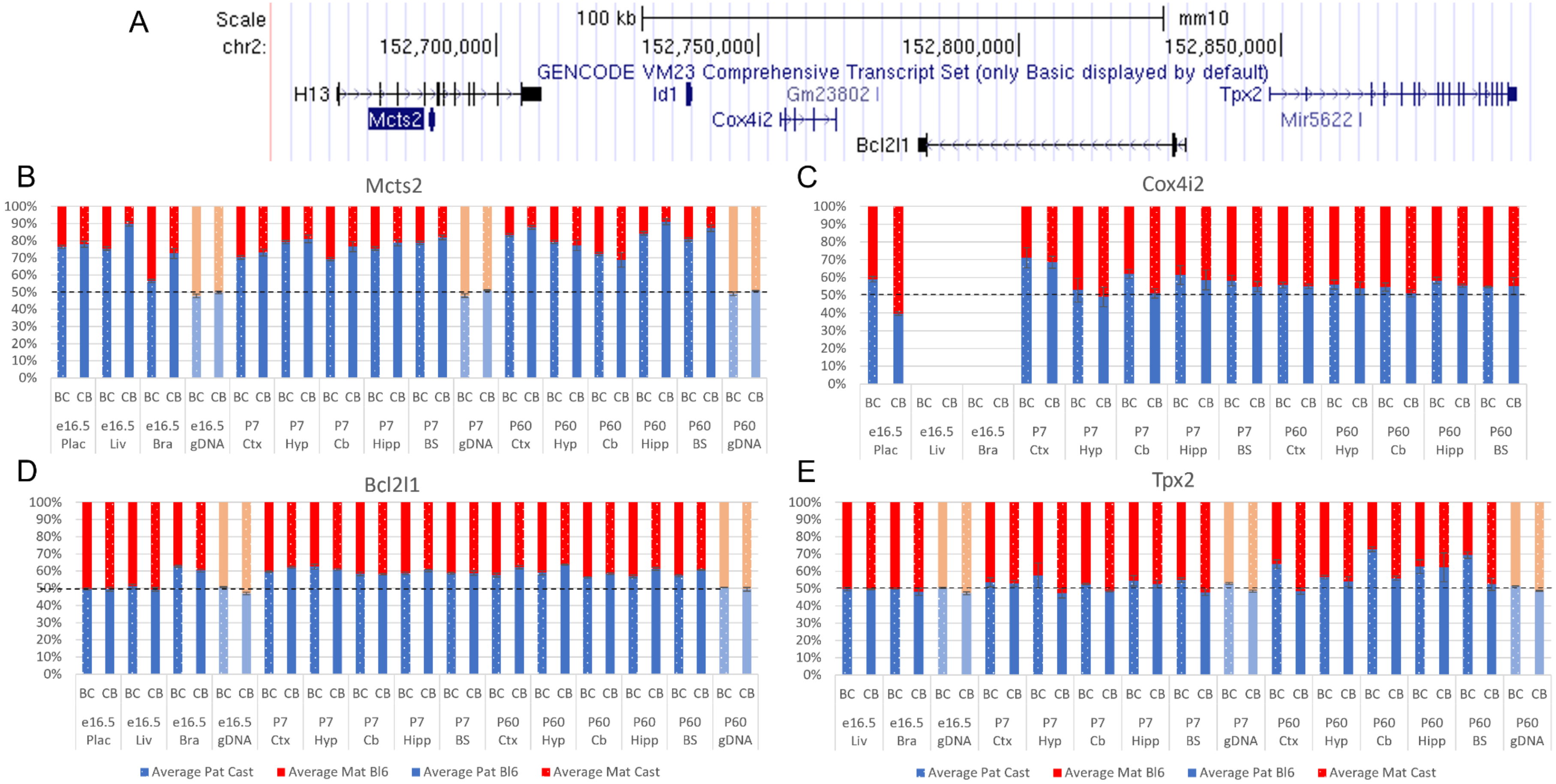


Figure S4 (A) *Mcts2* imprinted region. (B-E) Allelic bias (%) in *Mcts2* (A), *Cox4i2* (B), *Bcl2l1* (C) and *Tpx2* (D). Graphs show mean expression (%) from the paternal allele (deep blue) and maternal allele (red) C57BL/6 x CastEiJ (BC) and 4 CastEiJ x C57BL/6 (CB) crosses. Castaneus allele is denoted by spotted pattern. Standard error of the mean is shown, N = 3 or 4.

**Supplementary Figure S5**

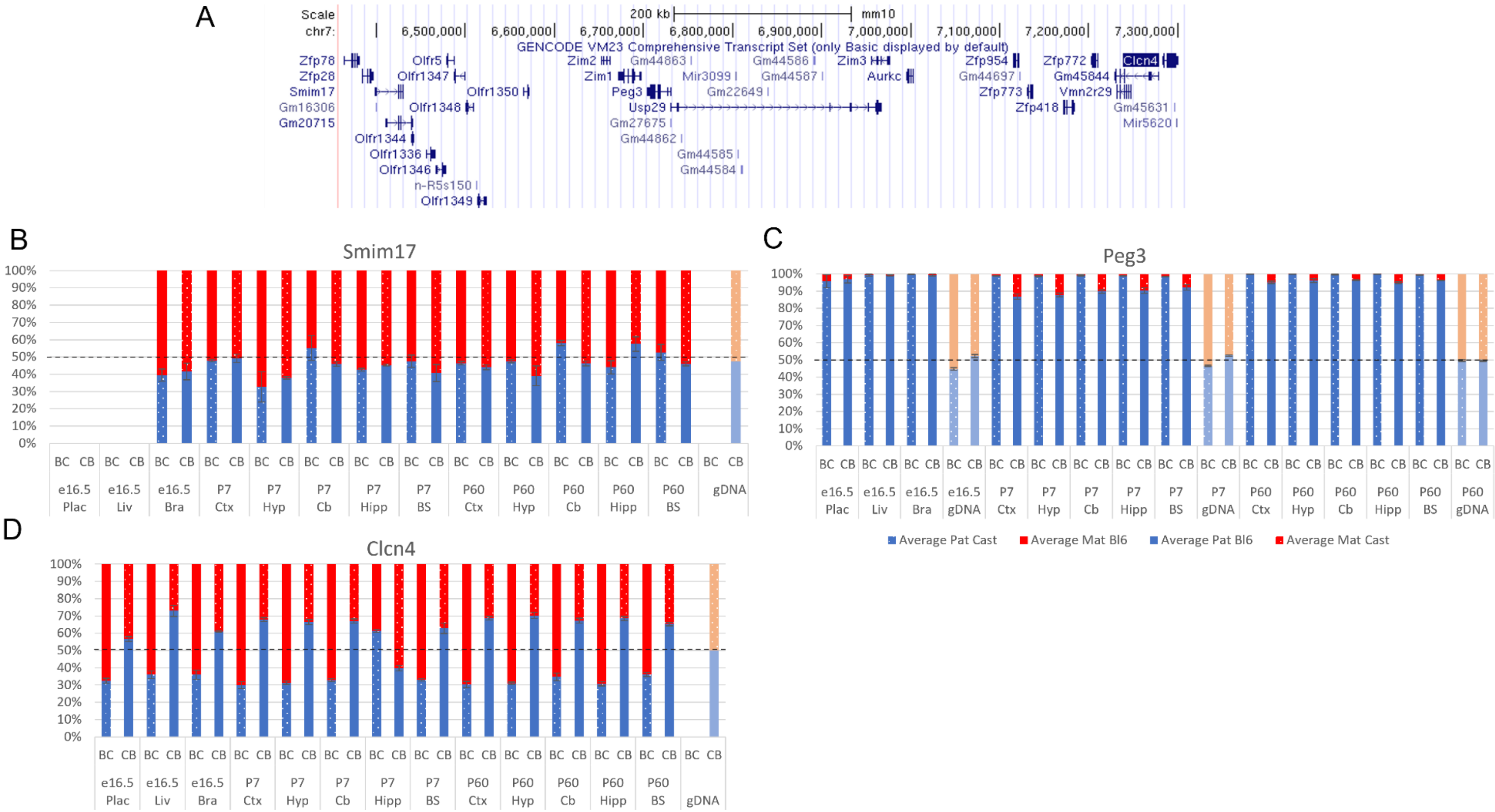


Figure S5 (A) *Peg3* imprinted region. (B-D) Allelic bias (%) in *Smim17* (A), *Peg3* (B) and *Clcn4* (C). Graphs show mean expression (%) from the paternal allele (deep blue) and maternal allele (red) C57BL/6 x CastEiJ (BC) and 4 CastEiJ x C57BL/6 (CB) crosses. Castaneus allele is denoted by spotted pattern. Standard error of the mean is shown, N = 3 or 4. *Clcn4* data are normalised for an amplification bias in gDNA.

**Supplementary Figure S6**

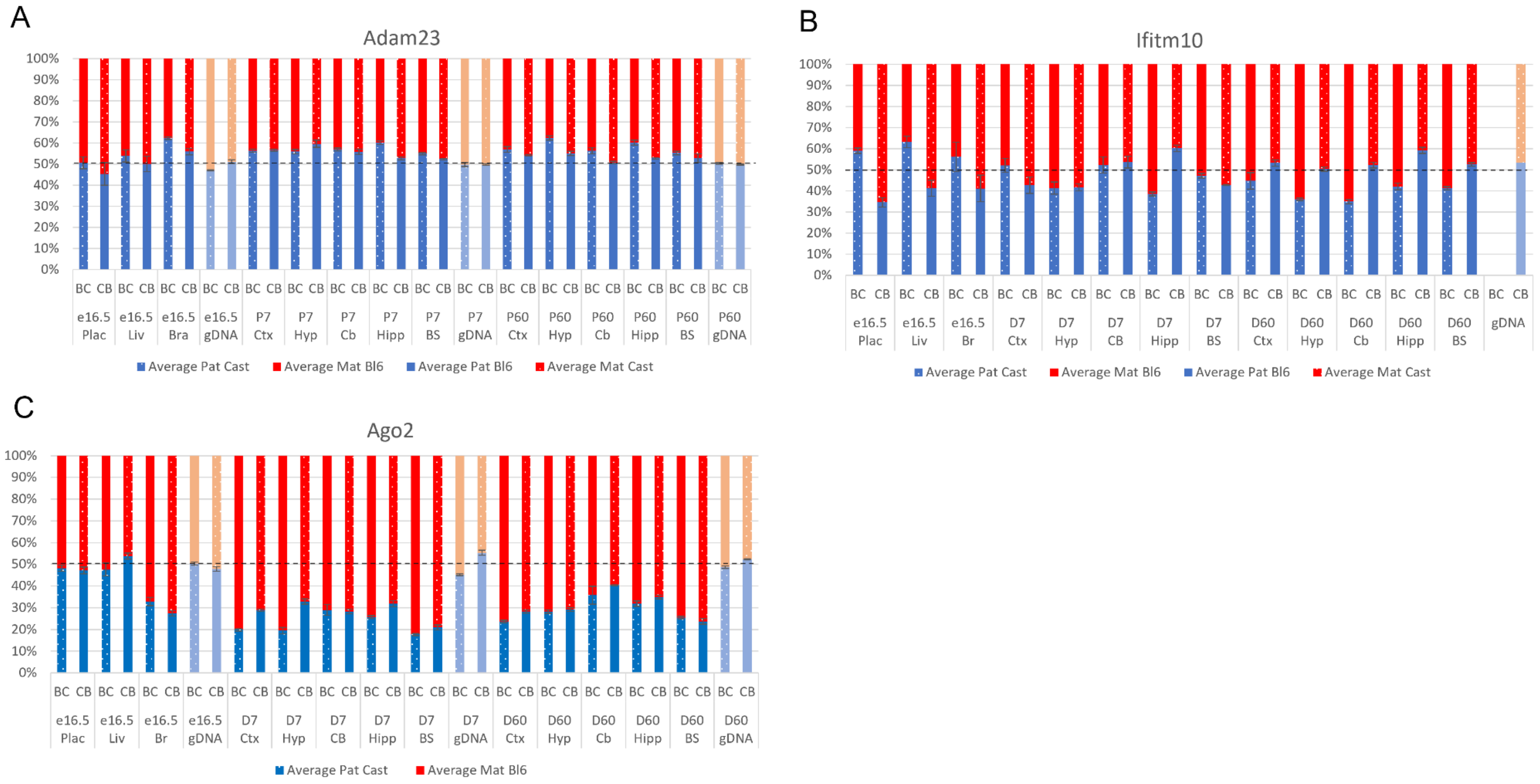


Figure S6 Allelic bias (%) in *Adam23* (A), *Ifitm10* (B) and *Ago2* (C). Graphs show mean expression (%) from the paternal allele (deep blue) and maternal allele (red) C57BL/6 x CastEiJ (BC) and 4 CastEiJ x C57BL/6 (CB) crosses. Castaneus allele is denoted by spotted pattern. Standard error of the mean is shown, N = 3 or 4. *Adam23* and *Iftim10* data are normalised for an amplification bias in gDNA.

**Supplementary Tables**

Table S1 – Study information for Babak et al. (2015), Bonthuis et al. (2015), Crowley et al. (2015) and Perez et al. (2015).

Table S2 – All genes called in original studies.

Table S3 – Overlapping Novel genes called in original studies.

Table S4 – All genes called in this study using ISoLDE.

Table S5 – Number of genes called in individual tissues in original study and this study.

Table S6 – Strain biased genes called in Dataset B and Dataset C in this study.

Table S7 – overlapping genes called in this study and the original one. This list includes genes generated in the undetermined list by the ISoLDE pipeline.

Table S8 – list of primers used in study.
